# Learning 2-in-1: Towards Integrated EEG-fMRI-Neurofeedback

**DOI:** 10.1101/397729

**Authors:** Lorraine Perronnet, Anatole Lécuyer, Marsel Mano, Mathis Fleury, Giulia Lioi, Claire Cury, Maureen Clerc, Fabien Lotte, Christian Barillot

**Affiliations:** University of Rennes, CNRS, Inria, Inserm, IRISA UMR 6074, Empenn Team ERL U 1228, Rennes, France; University of Rennes, CNRS, Inria, IRISA UMR 6074, Hybrid Project Team, Rennes, France; Inria, Athena Project Team, Sophia Antipolis, France; Inria, Potioc Project Team, Bordeaux, France; LaBRI (Univ. Bordeaux / CNRS / Bordeaux INP), Bordeaux, France

## Abstract

Neurofeedback (NF) allows to exert self-regulation over specific aspects of one’s own brain activity by returning information extracted in real-time from brain activity measures. These measures are usually acquired from a single modality, most commonly electroencephalography (EEG) or functional magnetic resonance imaging (fMRI). EEG-fMRI-neurofeedback (EEG-fMRI-NF) is a new approach that consists in providing a NF based simultaneously on EEG and fMRI signals. By exploiting the complementarity of these two modalities, EEG-fMRI-NF opens a new spectrum of possibilities for defining bimodal NF targets that could be more robust, flexible and effective than unimodal ones. Since EEG-fMRI-NF allows for a richer amount of information to be fed back, the question arises of how to represent the EEG and fMRI features simultaneously in order to allow the subject to achieve better self-regulation. In this work, we propose to represent EEG and fMRI features in a single bimodal feedback (integrated feedback). We introduce two integrated feedback strategies for EEG-fMRI-NF and compare their early effects on a motor imagery task with a between-group design. The BiDim group (n=10) was shown a two-dimensional (2D) feedback in which each dimension depicted the information from one modality. The UniDim group (n=10) was shown a one-dimensional (1D) feedback that integrated both types of information even further by merging them into one. Online fMRI activations were significantly higher in the UniDim group than in the BiDim group, which suggests that the 1D feedback is easier to control than the 2D feedback. However subjects from the BiDim group produced more specific BOLD activations with a notably stronger activation in the right superior parietal lobe (BiDim > UniDim, p < 0.001, uncorrected). These results suggest that the 2D feedback encourages subjects to explore their strategies to recruit more specific brain patterns. To summarize, our study shows that 1D and 2D integrated feedbacks are effective but also appear to be complementary and could therefore be used in a bimodal NF training program. Altogether, our study paves the way to novel integrated feedback strategies for the development of flexible and effective bimodal NF paradigms that fully exploits bimodal information and are adapted to clinical applications.

## 1 INTRODUCTION

Neurofeedback is the process of feeding back real-time information to an individual about his/her ongoing brain activity so that he/she can learn to control some aspects of it with potential for functional (behavioral, physical, cognitive, emotional) improvements (Sulzer et al., 2013; Thibault et al., 2015; Sitaram et al., 2016; Arns et al., 2017). NF has been investigated for a wide range of clinical (Birbaumer et al., 2009) and non clinical applications (Sulzer et al., 2013; Thibault et al., 2015, 2016; Sitaram et al., 2016; Arns et al., 2017; Gruzelier, 2014). However its effective deployment in the clinical practice is being held back by the debated evidence about its efficiency, most likely as a result of poor study design and lack of established guidelines and knowledge about the underlying mechanisms of NF (Thibault et al., 2016; Perronnet et al., 2016). In recent years, increasingly rigorous approaches are becoming the new standard (Ros et al., 2019; Sulzer et al., 2013; Stoeckel et al., 2014; Thibault et al., 2016), and new studies are digging into the mechanisms (Ninaus et al., 2013; Sitaram et al., 2016; Emmert et al., 2016; Birbaumer et al., 2013; Kober et al., 2013) as well as the methodological aspects of NF (Sorger et al., 2019; Emmert et al., 2017; Krause et al., 2017; Sorger et al., 2016; Sepulveda et al., 2016). However another reason for the debated efficiency of current approaches might be the inherent limitations of single imaging modalities (Biessmann et al., 2011; Fazli et al., 2015). Indeed, most NF approaches rely on the use of a single brain imaging modality such as EEG (Hammond, 2011), fMRI (Sulzer et al., 2013), functional near infra-red spectroscopy (fNIRS) (Mihara et al., 2012; Kober et al., 2014) or magnetoencephalography (Lal et al., 2005; Sudre et al., 2011; Buch et al., 2008). Each of these modalities is sensitive to a particular biophysical phenomenon related to the brain activity and comes with technical and physiological limitations (Biessmann et al., 2011). NF studies often report a significant proportion of subjects (usually about 30%) that are not able to self-regulate their brain activity (Alkoby et al., 2018). In the brain-computer-interface (BCI) community this phenomenon is known as BCI-deficiency and might originate from non optimal features, flaws from the design (Lotte et al., 2013; Chavarriaga et al., 2017), but also from anatomo-physiological factors that would make some subjects less responsive to certain modalities (Zich et al., 2015). EEG is the most popular NF modality for it is portable, non-invasive and benefits from millisecond temporal resolution. However, its spatial resolution is limited by volume conduction of the head and the number of electrodes. Also, source localization from EEG is inaccurate because of the ill-posed inverse problem (Grech et al., 2008; Baillet et al., 2001). fMRI is being increasingly used for NF as it allows to regulate even deeper brain regions with high spatial resolution (Sulzer et al., 2013). However its temporal resolution is limited by the time required to acquire one brain volume (hundreds of milliseconds), and the fact that the hemodynamic response peak is delayed of 4-6s from the neuronal onset and that it acts like a low-pass filter that smears out the neuronal response.

Bimodal EEG-fMRI-neurofeedback (EEG-fMRI-NF) is a new neurofeedback (NF) approach that consists of using information coming simultaneously from EEG and fMRI in real-time to allow the subject to regulate electrophysiological and hemodynamic activities of their brain at the same time (Zotev et al., 2014). The feasibility of this approach was demonstrated by Zotev et al. who hypothesized that it could be more efficient than the unimodal approaches (Zotev et al., 2014). Bimodal EEG-fMRI-NF was compared against unimodal EEG-NF and fMRI-NF in a recent study (Perronnet et al., 2017). This study suggested that EEG-fMRI-NF could indeed be more specific or more engaging than EEG-NF as demonstrated by higher BOLD activations during EEG-fMRI-NF than during EEG-NF. It also highligted that during bimodal EEG-fMRI-NF subjects could happen to regulate more one modality than the other suggesting the existence of specific mechanisms involved when learning to regulate simultaneously hemodynamic and electrophysiological aspects of the brain activity.

EEG and fMRI share mutual information yet also contain important distinct features. However their degree of overlap is hard to predict. In the context of NF, the information coming from EEG and fMRI could therefore benefit from being integrated in order to be used as an efficient feedback. Yet integrating EEG and fMRI data is a real challenge (Biessmann et al., 2011; Jorge et al., 2014; Fazli et al., 2015; Lahat et al., 2015). Multimodal data integration methods are categorized as asymmetrical (EEG-informed fMRI, fMRI-informed EEG) and symmetrical (data fusion, model-driven or data-driven) (Biessmann et al., 2011; Jorge et al., 2014; Lahat et al., 2015). For NF purpose the integration method should be applicable in real-time. As illustrated by Fig 1, the integration of multimodal data can theoretically occur at different levels: the raw measures level, the features level (high level or multivariate), the NF signal level or the feedback level (Fazli et al., 2015). It is also possible not to integrate EEG and fMRI data and simply show them as two separate feedbacks but we argue that this might be sub-optimal (see below). Integrating EEG and fMRI at the measures level in real-time does not seem feasible due to the considerable amount of information that it would represent. In hybrid BCI, output of different classifiers are usually passed to a meta-classifier (Fazli et al., 2012, 2015). In NF it is less common to use a classifier (Huster et al., 2014) and the feature often directly constitutes the NF signal. We argue that for EEG-fMRI-NF, EEG and fMRI integration can be done directly at the feedback level and that this already has strong implication.

**Figure 1.**
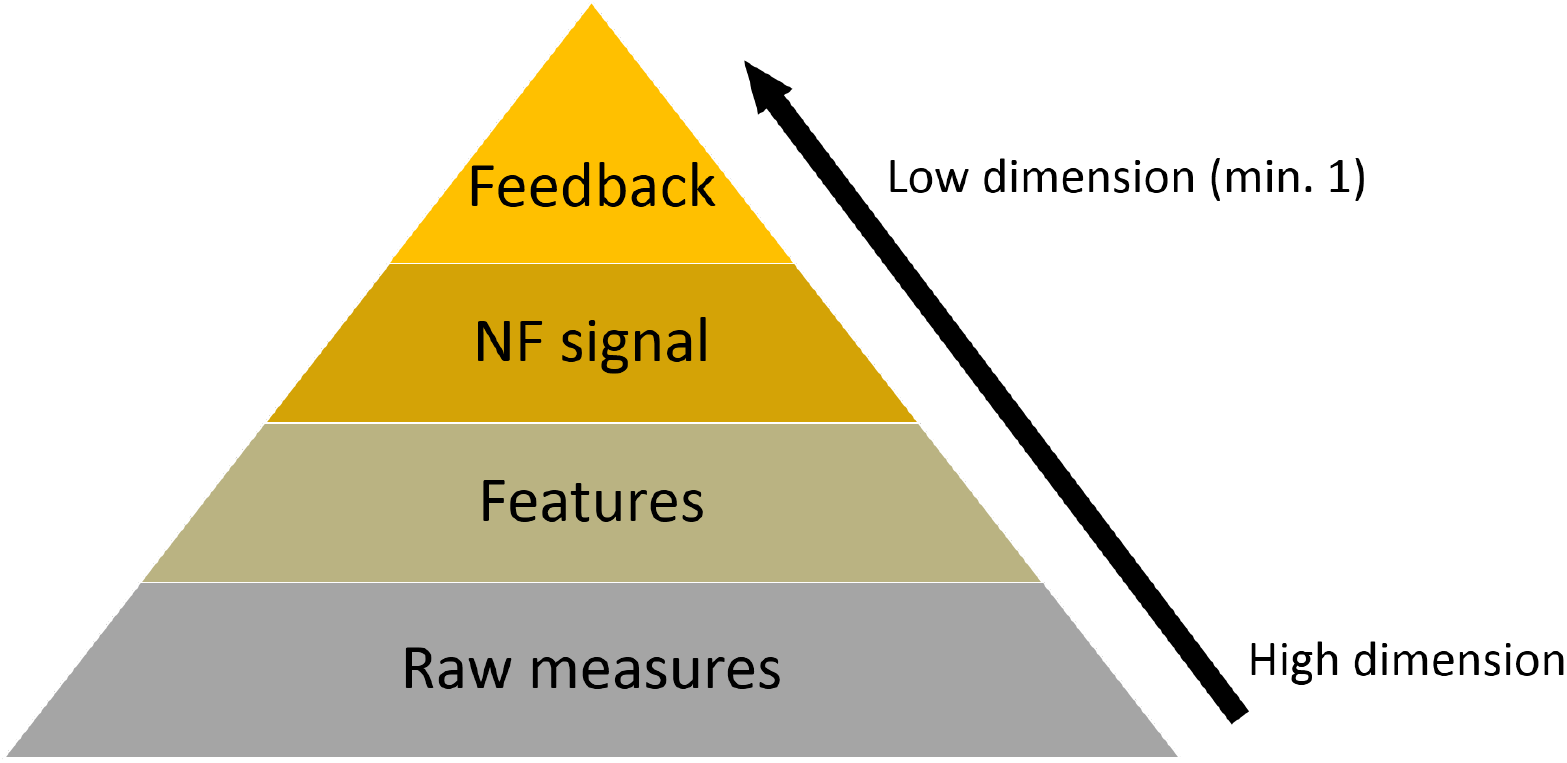
Possible levels of integration of EEG and fMRI information

Several unimodal NF/BCI studies have investigated the effects of feedback presentation as for example (Krause et al., 2017; Stoeckel et al., 2014; Sollfrank et al., 2016; Darvishi et al., 2017; Ono et al., 2014; Jeunet et al., 2015b; Kaufmann and Williamson, 2011) which is a central issue in NF/BCI design. One can refer to (Pillette, 2019) for a recent review feedback presentation in BCI. In the case of bimodal NF, feedback design might be even more critical as there is more information to display and as the EEG and fMRI information have different spatio-temporal dynamic properties. To our knowledge, no previous work has addressed the question of how to represent the EEG and fMRI information simultaneously and how the bimodal representation would affect NF performance.

In their pioneering work Zotev et al. naturally extended the classical thermometer feedback to bimodal NF by juxtaposing two feedback gauges, one for EEG and one for fMRI (Zotev et al., 2014). Though this has the advantage of clearly and fully representing both features, this could suffer from a few drawbacks. Firstly, it might be suboptimal in term of cognitive load (Sweller et al., 1998; Gaume et al., 2016) because the subject has to concentrate on two gauges. Secondly, the fact that the representations of both signals are separated seem to imply that there are two targets to reach. Therefore the regulation task might be perceived by the subject as two simultaneous regulation tasks instead of one. Thirdly, it can be misleading when the subject tries to interpret how both features evolve in time, especially when they exhibit inconsistencies. Lastly, it does not fully exploit the possibility of using a NF target defined by the state of both features.

In contrast to representing the EEG and fMRI features with two separate feedbacks, we propose an *integrated* feedback: that is a single feedback having a single NF target characterized by the state of both features. This type of feedback has the advantage of encouraging the subject to perceive the bimodal NF task as a single self-regulation task and offers great flexibility in the definition of bimodal NF targets. In this study, we introduce two integrated feedback strategies (illustrated in 2) and compare their early effects on an EEG-fMRI-NF guided motor-imagery task with a between-group design in order to evaluate which strategy is better than the other on a single EEG-fMRI-NF session. The performance of each strategy is evaluated in terms of sensitivity (activation level) and spatial specificity of the motor-imagery related EEG and fMRI activations.

## 2 MATERIAL AND METHODS

The study was conducted at the Neurinfo platform (CHU Pontchaillou, Rennes, France) and was approved by the Institutional Review Board (OSS-IRM, NCT number: NCT03440983). Twenty right-handed NF-naive healthy volunteers with no prior MI-NF experience (mean age: 35 *±* 10.6 years, 10 females) participated in the study. Participants were randomly assigned to the bi-dimensional (BiDim; mean age: 37 *±* 14 years, 5 females) or to the uni-dimensional (UniDim; mean age: 33 *±* 6.2 years, 5 females) group. Throughout the whole experiment, the participants were lying down in the MR bore and wearing a 64 channel MR-compatible EEG cap. The first integrated feedback strategy is a *two-dimensional* (2D) plot in which each dimension depicts the information from one modality. The second integrated feedback strategy is a *one-dimensional* (1D) gauge that merges both information into one and therefore has a higher degree of integration than the 2D feedback.

### 2.1 Hypotheses

Figure 2 illustrates the separate feedback (adapted from (Zotev et al., 2014)) as well as the integrated 2D and 1D feedback strategies and summarizes their potential advantages and drawbacks. The proposed integrated 2D feedback consists of a ball moving on a 2D diamond-shape plot, the left dimension representing the EEG feature and the right dimension representing the fMRI feature. This type of feedback was introduced in our previous work (Perronnet et al., 2017) and we propose here an upgraded version in which the plot background delineates regions that indicate prefered direction of effort, encouraging the subject to regulate EEG and fMRI equitably. Our integrated 1D feedback consists of a ball moving in a gauge, the ball position representing the average of the EEG and fMRI features. The background of the 1D gauge is splitted into four regions to give the subject reference points. Interestingly, the separate feedback and the integrated 2D feedback strategies share common advantages and drawbacks because they both map the bimodal information onto two dimensions. On the good side, they fully represent the EEG and fMRI features and allow to discriminate between both, but as a counterpart the difference of update rates is perceivable and inconsistencies between the two features can be misleading. The integrated 2D feedback is visually more optimal than the separate feedback as subjects only need to look at a single representation. However it might be complex to apprehend as the subjects need to understand how the ball travels in the 2D space. Subjects would therefore probably need more time to get used to this feedback. But the information it conveys is highly meaningful regarding how the subject is regulating both features at the same time. When the ball is on one side it means that the subject is controlling more one feature than the other, but when the ball is on the diagonal it means that the subject is controlling both features equally. The integrated 1D feedback has the advantage of being simpler to understand but as a counterpart it is less informative than the two other NF metaphors.

**Figure 2.**
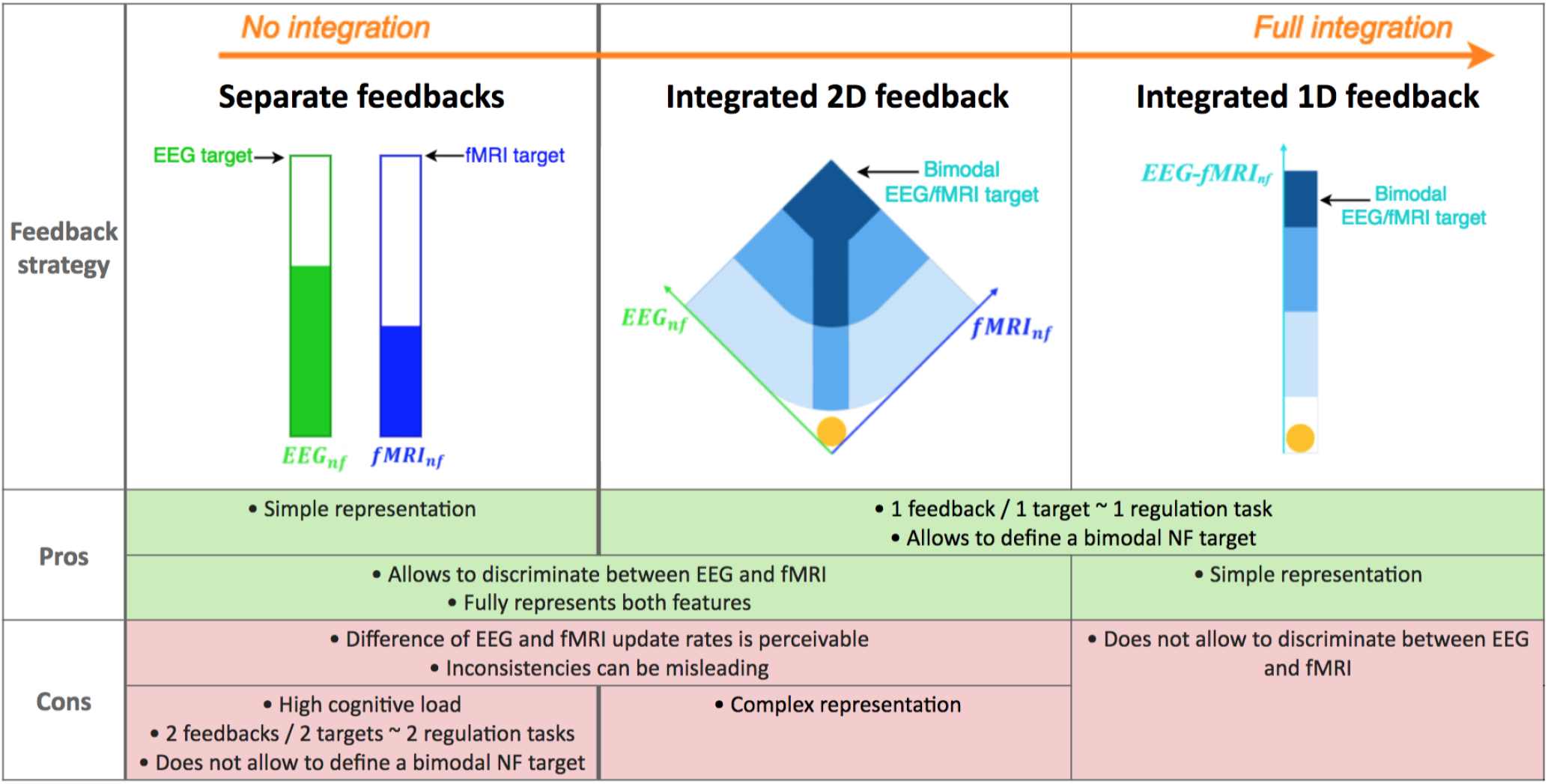
Summary of potential advantages and drawbacks of different bimodal feedback strategies: separate, integrated 2D, integrated 1D. As opposed to the separate feedback strategy proposed by pioneer authors in (Zotev et al., 2014), we introduce two novel integrated feedback strategies for bimodal EEG-fMRI-NF. The integrated 2D feedback consists of a ball moving in two dimensions, the left dimension representing the EEG feature and the right dimension representing the fMRI feature. Subjects in the BiDim group are shown the integrated 2D feedback. The integrated 1D feedback consists of a ball moving in one dimension, the ball position representing the average of the EEG and the fMRI features. Subjects in the UniDim group are shown the integrated 1D feedback. NB: in this study, we compare the integrated 2D and the integrated 1D strategies, however we do not evaluate integrated feedback strategies against the separate feedback strategy.

### 2.2 Experimental protocol

After signing an informed consent form describing the MR environment, the participants were verbally informed about the goal of both the study and the protocol. They were instructed that during the NF runs, they would be presented with a ball moving in two dimensions (for the BiDim group) or in a one-dimensional gauge (for the UniDim group) according to the activity in their motor regions as measured with EEG and fMRI (see Fig 2). Participants were told that they would have to bring the ball closer to the darker blue areas by imagining clenching their right-hand. This instruction was reminded in written form on the screen at the beginning of each NF run. More specifically we explained the participants that they would need to perform kinesthetic motor imagery (kMI) (Neuper et al., 2005) of their right-hand in order to control the ball. Kinesthetic motor imagery was defined as trying to feel the sensation of the motion rather than only visualizing it. We told them that they should try to control both dimensions, i.e. try to move the ball near the diagonal. These instructions were given verbally at the beginning of the experiment and reminded later if the participant asked for it. Participants were asked not to move at all, especially during the course of a run. Video monitoring of the inside of the MR tube allowed to check for whole-body movements of the participant.

After receiving the instructions and having the EEG cap setup, the participant was installed in the MR tube. The experimental protocol then consisted of: a structural 3D T1; a preliminary MI run without NF (MI pre), used to calibrate the NF targets (see section 2.5); three NF runs with a one minute break in between each; a post MI run without NF (MI post). The five EEG-fMRI functional runs employed a block-design alternating 8 times 20s of rest and 20s of task (see Figure 3).

**Figure 3.**
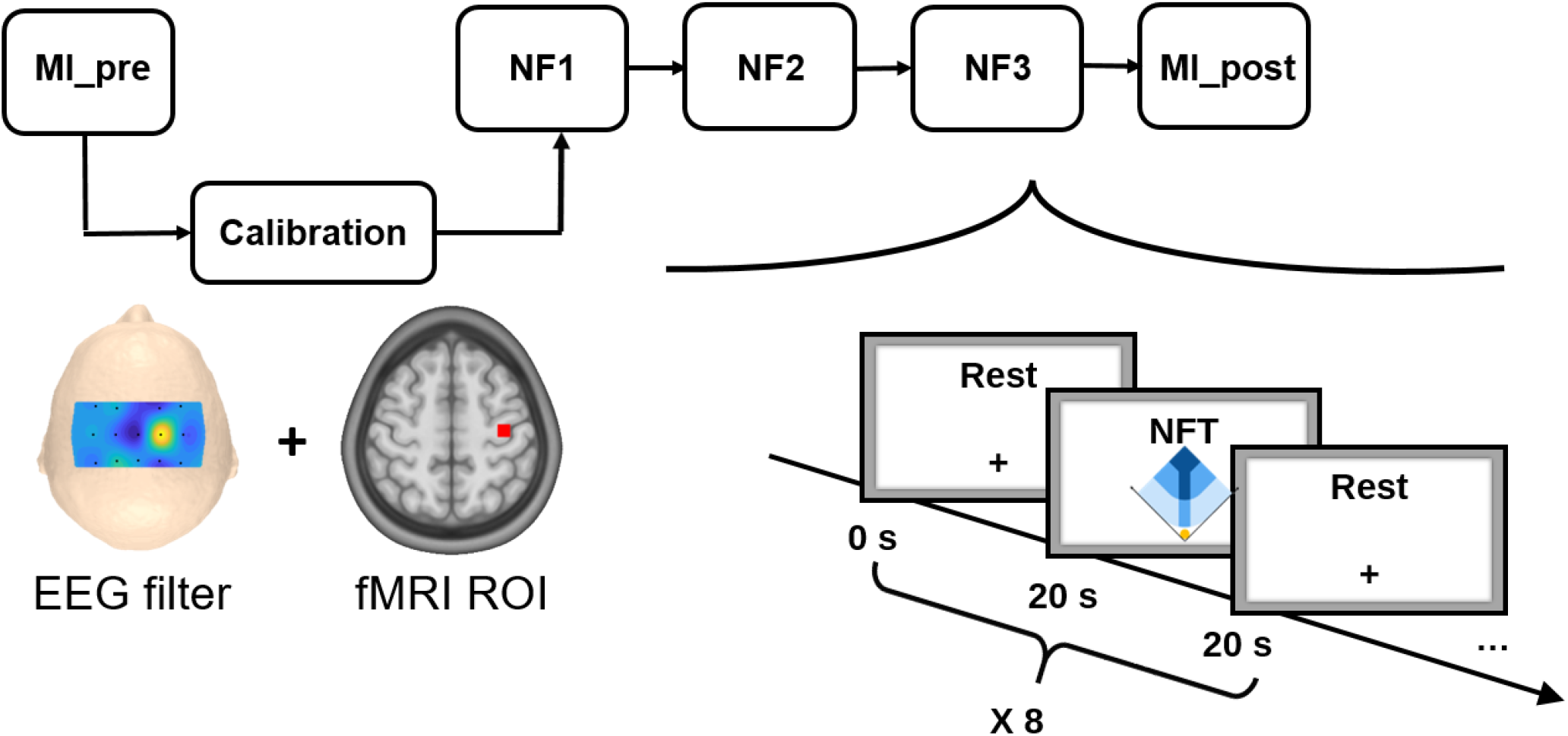
The experimental protocol consisted of 5 EEG-fMRI runs: a preliminary motor imagery run without NF (MI pre) used for calibration, three NF runs (NF1, NF2, NF3), and a post motor imagery run without NF (MI post). Each run consisted of a block design alternating 8 times 20s of rest and 20s of task.

During rest, the screen displayed a white cross and participants were asked to concentrate on the cross and not on the passed or upcoming task block. During task, the screen displayed the cue “Imagine right” as well as the feedback during NF runs. The feedback consisted of a yellow ball moving in a two-dimensional plot for the BiDim group or in a one-dimensional gauge for the UniDim group. The participants were instructed to bring the ball closer to the darker blue area by performing kinesthetic motor imagery of their right hand clenching. The EEG feature was defined as the event-related desynchronization (ERD) (Pfurtscheller and Lopes da Silva, 1999) in the [8-30Hz] band of the EEG data filtered with a subject specific spatial filter (see Section 2.5 and 2.4) and was updated every 250ms. The fMRI feature was defined as the mean BOLD in a subject-specific motor region-of-interest (ROI) (see Sections 2.5 and 2.4) and was updated at every repetition time (TR=1s). For the UniDim group, the ball position was the average of the EEG and fMRI features (*EEG_nf_* + *f MRI_nf_*)*/*2. For the BiDim group, the right axis depicted the normalized fMRI feature while the left axis depicted the normalized EEG feature. At the end of the experiment, the participants were asked to fill out a home-made questionnaire about their perceived performance motivation, fatigue, interest and difficulty in performing the NF task.

### 2.3 Data acquisition

EEG and fMRI data were simultaneously acquired with a 64-channel MR-compatible EEG solution from Brain Products (Brain Products GmbH, Gilching, Germany) and a 3T Verio Siemens scanner (VB17) with a 12 channel head coil. Foam pads were used to restrict head motion. EEG data was sampled at 5kHz with FCz as the reference electrode and AFz as the ground electrode. fMRI acquisition was performed using echo-planar imaging (EPI) with the following parameters: repetition time (TR) / echo time (TE) = 1000/23ms, FOV = 210 *×* 210*mm*^2^, voxel size = 2 *×* 2 *×* 4*mm*^3^, matrix size = 105 *×* 105, 16 slices, flip angle = 90^*°*^. Visual instructions and feedback were transmitted using the NordicNeurolab hardware and presented to the participant via an LCD screen and a rear-facing mirror fixed on the coil. As a structural reference for the fMRI analysis, a high resolution 3D T1 MPRAGE sequence was acquired with the following parameters: TR/TI/TE = 1900/900/2.26ms, GRAPPA 2, FOV = 256 *×* 256*mm*^2^ and 176 slabs, voxel size = 1 *×* 1 *×* 1*mm*^3^, flip angle = 90^*°*^. Our multimodal EEG/fMRI-NF system (Mano et al. 2017) integrates EEG and fMRI data streams via a TCP/IP socket. The EEG data is pre-processed with BrainVision Recview (Brain Products GmbH, Gilching, Germany) software for gradient and ballistocardiogram (BCG) artifact correction (see Section 2.4) and sent to Matlab (The MathWorks, Inc., Natick, Massachussets, United States) for further processing. The fMRI data is pre-processed online for slice-time correction and motion correction with custom Matlab code adapted from SPM8 (FIL, Wellcome Trust Centre for Neuroimaging, UCL, London, UK). EEG and fMRI NF features are then computed and translated as feedback with Psychtoolbox (Kleiner et al., 2007).

### 2.4 Real-time data processing

During NF runs, online gradient artifact correction and BCG correction of the EEG data were done in BrainVision Recview (Brain Products GmbH, Gilching, Germany) software. The gradient artifact correction in Recview is based on the average artifact subtraction (AAS) method (Allen et al., 2000). At the beginning and throughout the length of each experiment, we checked that the signal quality of the ECG channel was good because BCG artifact correction relies on the quality of the ECG channel. We would also reset the template regularly before the start of each run. We used an artifact subtraction template of 2000ms and 4 templates for template drift correction. The data was then down-sampled to 200Hz and low pass filtered at 50 Hz (48 db slope) with a Butterworth filter (order 8). The data were subsequently corrected for BCG artifact (Allen et al., 1998). The pulse model was searched in the first 15 seconds of the data. The pulse detection was based on a moving template matching approach with minimal pulse period of 800ms, minimum correlation threshold of 0.7, and amplitude ratio range from 0.6 to 1.2 relative to the pulse model. For pulse correction, a moving template was computed by averaging the 10 previously detected pulses, and the correction was done on a window length of [-100ms, 700ms] relatively to the R-peak. This corrected data was then sent to Matlab for feature extraction. The corrected data was filtered with the subject specific spatial filter computed during the calibration phase (see Section 2.5). The band power in the [8-30Hz] band was then computed on this filtered data using the periodogram and a 2s window size, and it was normalized with the following ERD-like (Pfurtscheller and Lopes da Silva, 1999) formula:

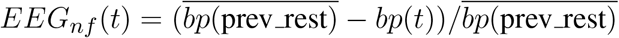

where *bp*(*t*) is the power at time t, 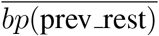 is the average power over the previous rest block (values between the fourteen and the nineteen seconds). Finally, the EEG features were smoothed over the last four values translated as visual feedback every 250ms and normalised. EEG features were normalised by dividing it with the value reached 30% of the time by the subject during calibration phase.

The fMRI signal was pre-processed online for motion correction, slice-time correction and then the fMRI NF feature was computed according to the following definition:

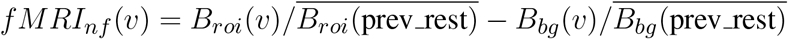

where *B_roi_*(*v*) (respectively *B_bg_*(*v*)) is the average BOLD signal in the ROI (respectively in the background (BG)) at volume v, and 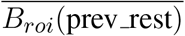 (respectively 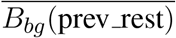 is the ROI (respectively BG) baseline obtained by averaging the signal in the ROI (respectively in the BG) from the fourteenth to the nineteenth second (to account for the hemodynamic delay) of the previous rest block. The background was defined as a large slice (slice 6 out of 16) in deeper regions and used to cancel out global changes. Finally the fMRI feature was smoothed over the last three volumes, normalised by the value reached 30% of the time by the subject during its calibration phase, and translated as visual feedback every second.

### 2.5 Calibration

In order to define subject-specific NF features, right at the end of the MI pre run, the MI pre EEG and fMRI data were pre-processed and analyzed to extract a spatial filter for EEG NF features, a BOLD ROI for fMRI NF features and to estimate the corresponding 30%-reaching threshold values for NF normalisation.

#### 2.5.1 EEG calibration

At the end of the MI pre run, the MI pre data was pre-processed similarly to what was done in real-time (see Section 2.4) except that the BCG correction was done semi-automatically. Using the Common Spatial Pattern (CSP) method (Ramoser et al., 2000), we then computed the pair of spatial filters that best maximized the difference in [8-30Hz] power between rest and task blocks on 18 channels located over the motor regions (C3, C4, FC1, FC2, CP1, CP2, FC5, FC6, CP5, CP6, C1, C2, FC3, FC4, CP3, CP4, C5, C6). In case the CSP filter did not correlate with the task during MI pre, we used a laplacian filter over C3 instead (Nunez et al., 1997).

#### 2.5.2 fMRI calibration

MI pre fMRI data was pre-processed for slice-time correction, spatial realignment and spatial smoothing with a 6mm Gaussian kernel with SPM8. A first-level general linear model (GLM) analysis was then performed. The fMRI ROI was defined by taking a 9 *×* 9 *×* 3 box around the maximum of activation (constrained to the left motor area) of the thresholded T-map (*task > rest*, *p <* 0.001, *k >* 10). The fMRI feature was then computed on this MI pre data (see Section 2.4).

### 2.6 Offline analysis

#### 2.6.1 EEG analysis

For offline analysis, EEG signal was pre-processed similarly to what was done in real-time (see Section 2.4) except that the BCG correction was done semi-automatically. For each subject and run, we checked that the ECG peaks were well detected by adapting the parameters and by correcting or selecting the ECG peaks manually when necessary.

To analyze how the participants regulated their EEG NF feature, we re-computed the ERD values on offline pre-processed data filtered with the online spatial filter as defined in 2.4 except that the baseline was not computed sliding-block-wise, but instead by averaging power values after the first second and before the nineteenth second of all rest blocks. We refer to this feature as “online EEG NF”.

As the amount of calibration data was limited and as participants had no prior MI training, it is possible that the filter from the calibration was suboptimal. Therefore we also extracted the EEG NF values on data filtered with a posthoc spatial filter. We refer to this feature as “posthoc EEG NF”. The posthoc spatial filter was computed as for the online setup (see Section 2.5) except that it was computed on the concatenation of MI pre, NF1, NF1 and NF3 instead of MI pre only.

For statistical analysis, the EEG NF values were standardized to z-scores by considering for each subject their mean and standard deviation over MI pre, NF1, NF2, NF3, MI post. For each run the standardized EEG NF values were averaged by considering the values between the first and the nineteenth second of all NF blocks but the first. The mean EEG NF over NF1, NF2 and NF3 was averaged to get the mean EEG NF 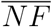. We also considered max *NF_i_* as the best mean EEG NF over the three NF runs. We refer to the best NF run regarding the EEG feature as *maxNF_eeg_*.

#### 2.6.2 fMRI analysis

The fMRI data from each of the five runs (MI pre, NF1, NF2, NF3, MI post) was pre-processed and analyzed with AutoMRI (Maumet, 2013), a proprietary software for fMRI analysis based on SPM8. Pre-processing included slice-time correction, spatial realignment, co-registration to the 3D T1, followed by spatial smoothing with a 8 mm Gaussian kernel. A first-level and second-level general linear model (GLM) analysis was performed. The first-level GLM included the canonical HRF for the task as well as its temporal and dispersion derivatives. For the second-level GLM analysis, the individual data were normalized to the Montreal Neurological Institute (MNI) template and grouped using a mixed effects linear model. The activation maps were corrected for multiple comparisons using Family-Wise error (FWE) correction (*p <* 0.05 with cluster size > 10 voxels).

To analyze how the participants regulated the BOLD signal in the *online ROI*, we extracted the ROI percent signal change (PSC) on offline p re-processed d ata. F or e ach p articipant a nd e ach r un, the registered fMRI values were high-pass filtered (100 seconds) to remove the linear drift, averaged in the online ROI and transformed into PSC using the formulae 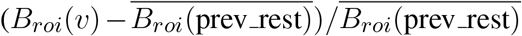. We refer to this feature as “online fMRI NF”.

Because NF training affects patterns beyond the one being fed back (Wander et al., 2013; Kopel et al., 2016), the same procedure was done to extract the PSC in a *posthoc ROI* defined by computing individually an average activation map over NF1, NF2 and NF3 and taking a 9 *×* 9 *×* 3 box around the maximum of activation (constrained to the left motor area). We refer to this feature as “posthoc fMRI NF”. Finally the posthoc fMRI NF values were standardized to z-scores by considering for each subject their mean and standard deviation over MI pre, NF1, NF2, NF3, MI post. For each run the standardized posthoc fMRI NF values were averaged across the last 16 volumes of all NF blocks but the first. The mean PSC over NF1, NF2 and NF3 was averaged to get the mean NF PSC 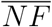. We also considered max *NF_i_* the best mean fMRI NF over the three NF runs. We refer to the best NF run regarding the fMRI feature as *maxNF_fmri_*.

#### 2.6.3 Statistical analysis

For each group (UniDim/BiDim), each modality (EEG/fMRI) and level of feature (online/posthoc) we conducted non-parametric Friedman tests of the differences among MI pre, 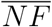, MI post, as well as Wilcoxon signed-rank tests (signrank Matlab function) between 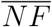 and MI pre as well as between max *NF_i_* and MI pre with Bonferroni correction (corrected p-value: 0.05/3 conditions = 0.0167). For between group comparison we computed a Wilcoxon test (ranksum Matlab function, equivalent to Mann-Whitney U-test) on 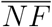. The tests were done both for the online fMRI NF and for the posthoc fMRI NF.

## 3 RESULTS

GLM analysis of both groups (UniDim + BiDim) revealed activations during NF (see Fig 4) in: bilateral premotor cortex (PMC) (BA 6) including left and right supplementary motor area (SMA), left and right inferior frontal gyrus (pars opercularis rolandic operculum) (BA 44), left and right inferior parietal lobule (IPL), left and right superior parietal lobule (SPL), left and right supramarginal lobule/gyrus (BA 40,BA 2,BA 48), left and right superior parietal (BA 7, BA 5), bilateral mid-cingulate cortex, left and right precuneus (BA 7). Deactivations were observed in right primary motor cortex (M1), left and right angular gyrus (BA 39), right cuneus (BA 18), left and right precuneus, left middle occipital (BA 10) and in the left inferior parietal lobule (BA 19).

**Figure 4.**
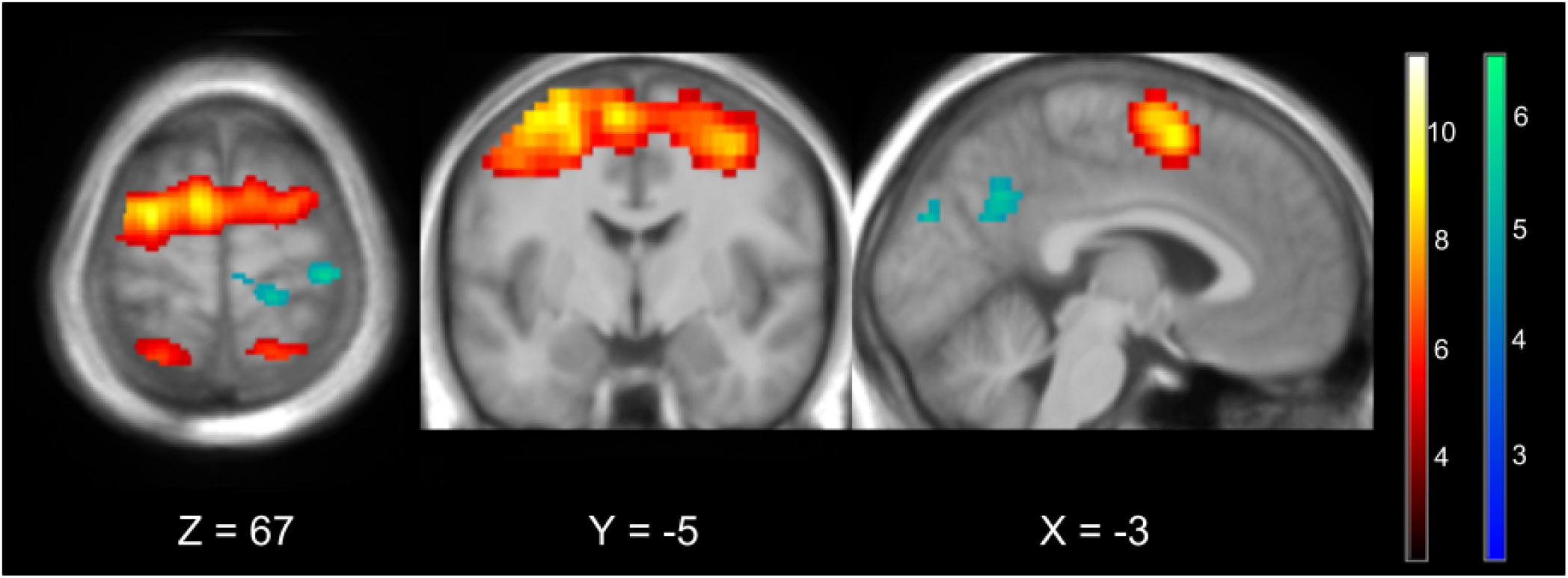
Average BOLD activations (in orange) and deactivations (in blue) over the three NF runs (NF1+NF2+NF3) in both groups (UniDim + BiDim) thresholded at p<0.05 FWE corrected

GLM analysis of the BiDim group during NF revealed activations in (Fig 5): Left PMC (BA 6) including SMA, left IPL (BA 40), left SPL (BA 7), right SPL (BA 5, BA 7), right superior occipital (BA 7). Deactivations were observed in right M1, (BA 4), left IPL (BA 19). GLM analysis of the UniDim group during NF revealed activations in (Fig 5): left and right PMC (BA 6) including left and right SMA, left IPL (BA 40), left superior parietal lobule (BA 40), left and right supramarginal lobule (BA 2). Deactivations were observed in the right angular gyrus (BA 39).

**Figure 5.**
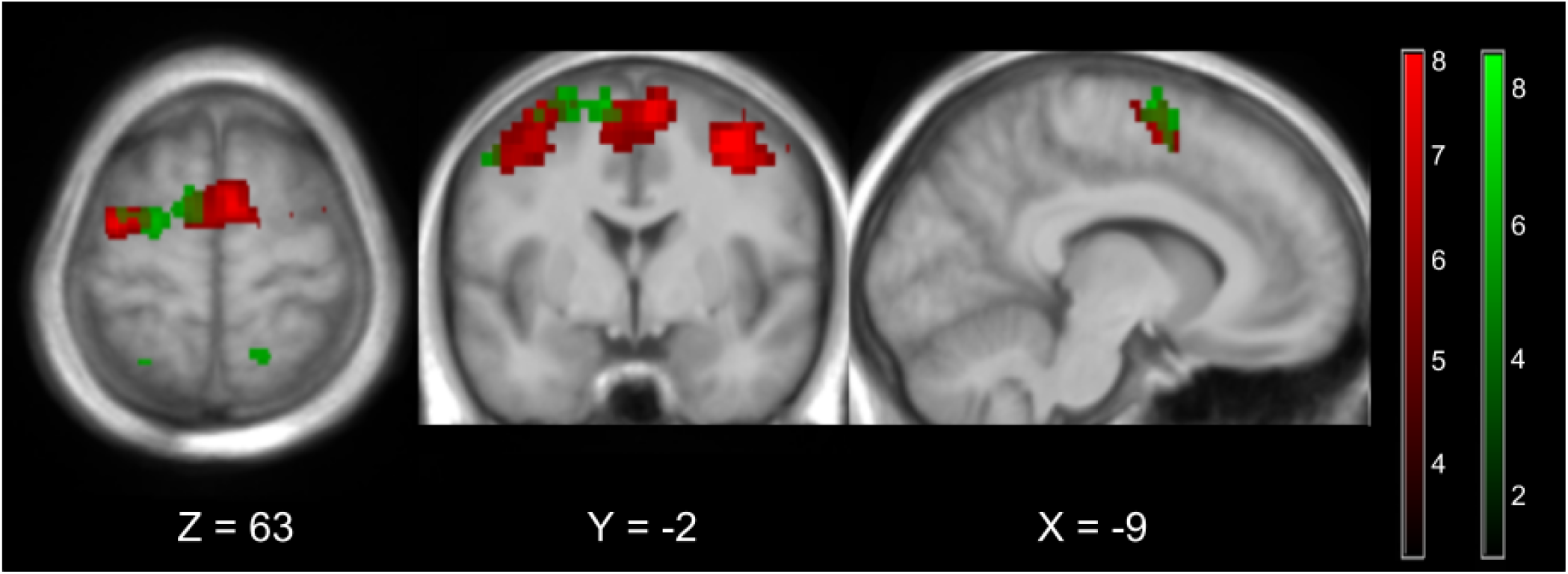
Average BOLD activations over the three NF runs (NF1+NF2+NF3) in each group thresholded at p<0.05 FWE corrected. Activations of the UniDim group are shown in red, and for the BiDim group in green.

**Figure 6.**
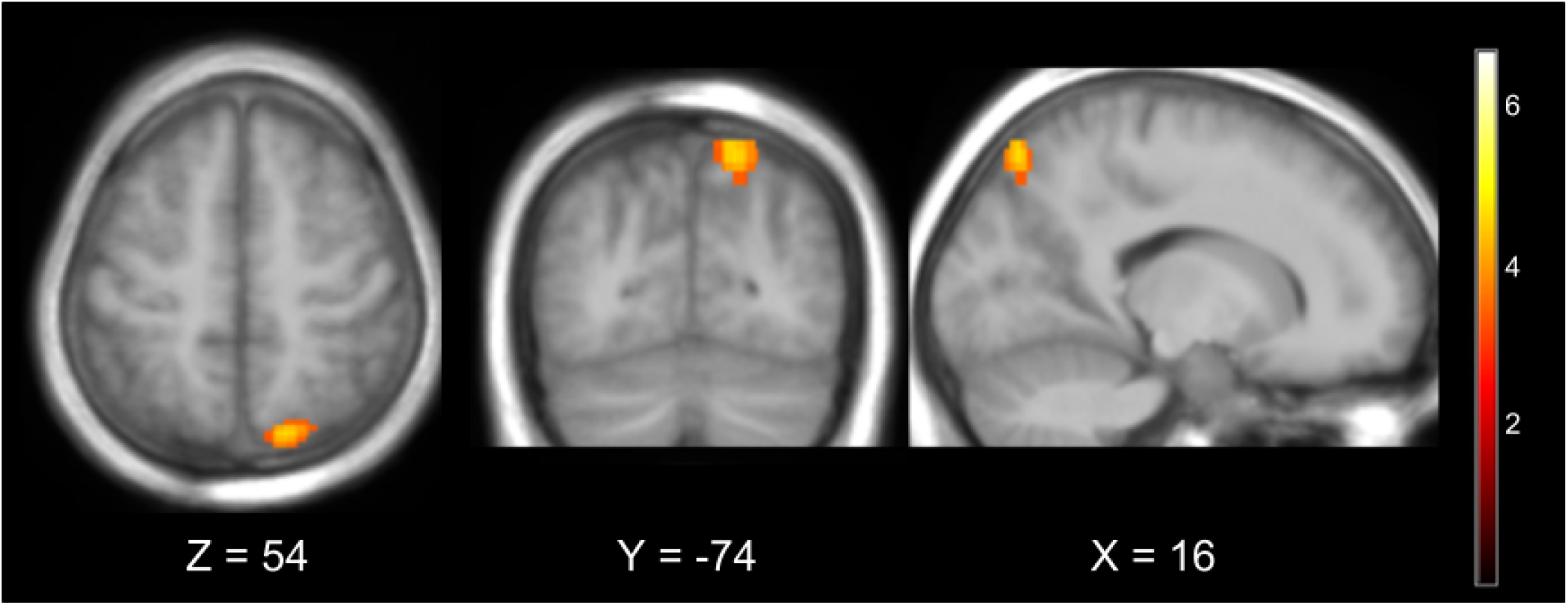
Group difference: BiDim>UniDim thresholded at p<0.001 uncorrected. The BiDim activated more the right superior parietal lobule (BA7).

**Figure 7.**
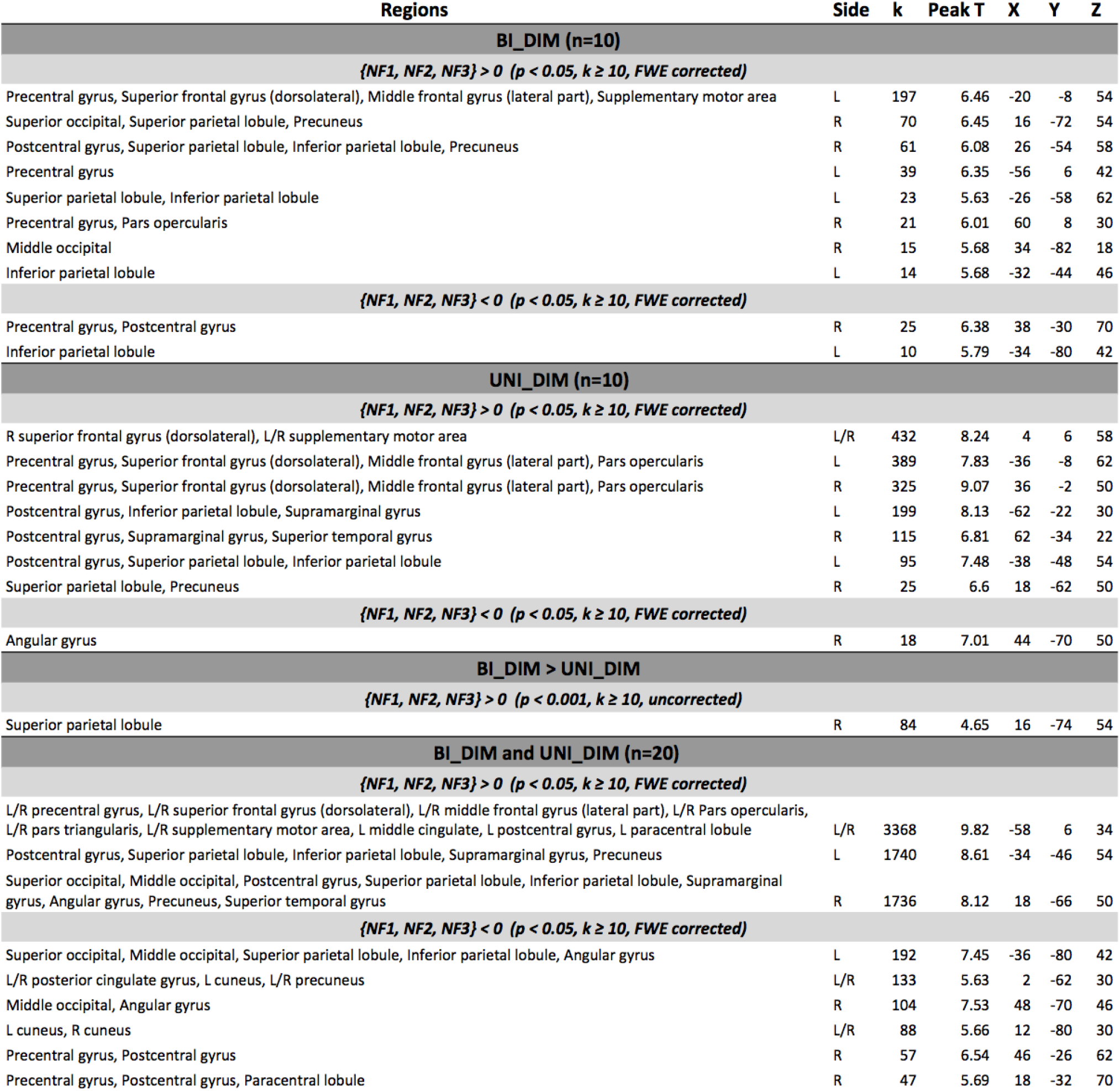
Anatomical labels, hemisphere, cluster size, peak t-value and MNI coordinates of significant group activation/deactivation clusters.

The BiDim group showed more activations (*p <* 0.001, uncorrected) than the UniDim group in the right superior parietal lobule (BA 7).

Friedman tests between MI pre, 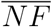 and MI post were significant for posthoc EEG in the BiDim group (p=0.045, *χ*^2^(2, 10) = 6.2) and for posthoc fMRI in the BiDim group (p=0.0136, *χ*^2^(2, 10)= 8.6).

Wilcoxon signed rank tests between MI pre and maxNF were significant for: online EEG (p=0.0098, signedrank = 52) and online fMRI (p=0.0195, signedrank = 50) in the UniDim group; posthoc EEG (p=0.0020, signedrank =55) and posthoc fMRI (p=0.0137, signedrank =51) in the UniDim group; and for posthoc EEG (p=0.0020, signedrank =55) and posthoc fMRI (p=0.0020, signedrank =55) in the BiDim group. Wilcoxon signed rank tests between MI pre and NF were significant for: posthoc EEG (p=0.0195, signedrank = 50) in the UniDim group; posthoc EEG (p=0.0273, signedrank = 49) and posthoc fMRI (p=0.0039, signedrank = 54) in the BiDim group. Results are summarized in Fig 8 and Fig 9. During the NF runs the fMRI NF in the online ROI was significantly higher in the UniDim group than in the BiDim group (Wilcoxon: z = 3.0615, ranksum = 146, p = 0.0022).

**Figure 8.**
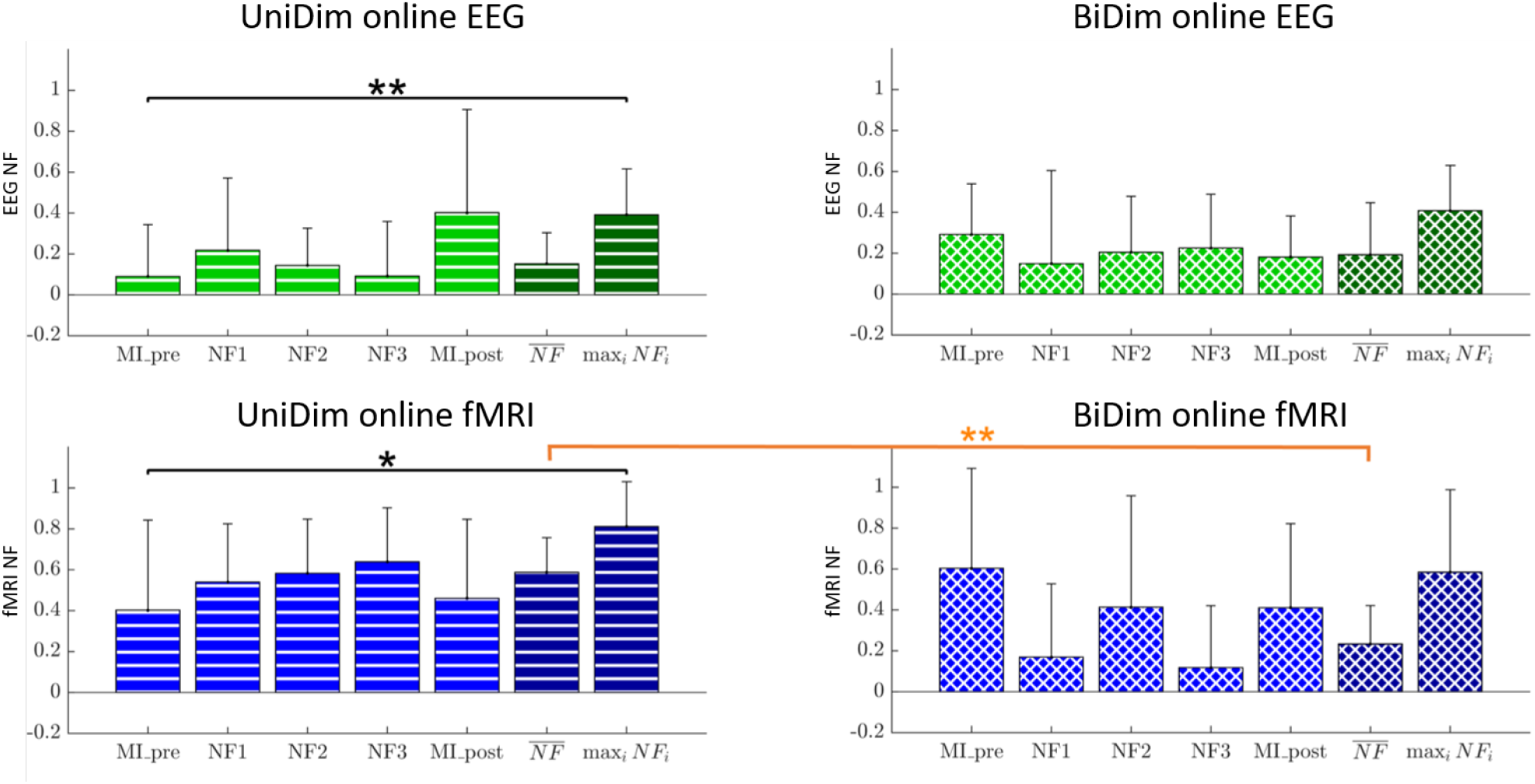
Group means (EEG/fMRI, online, z-scored) on each run with standard deviation + significance of Wilcoxon tests

**Figure 9.**
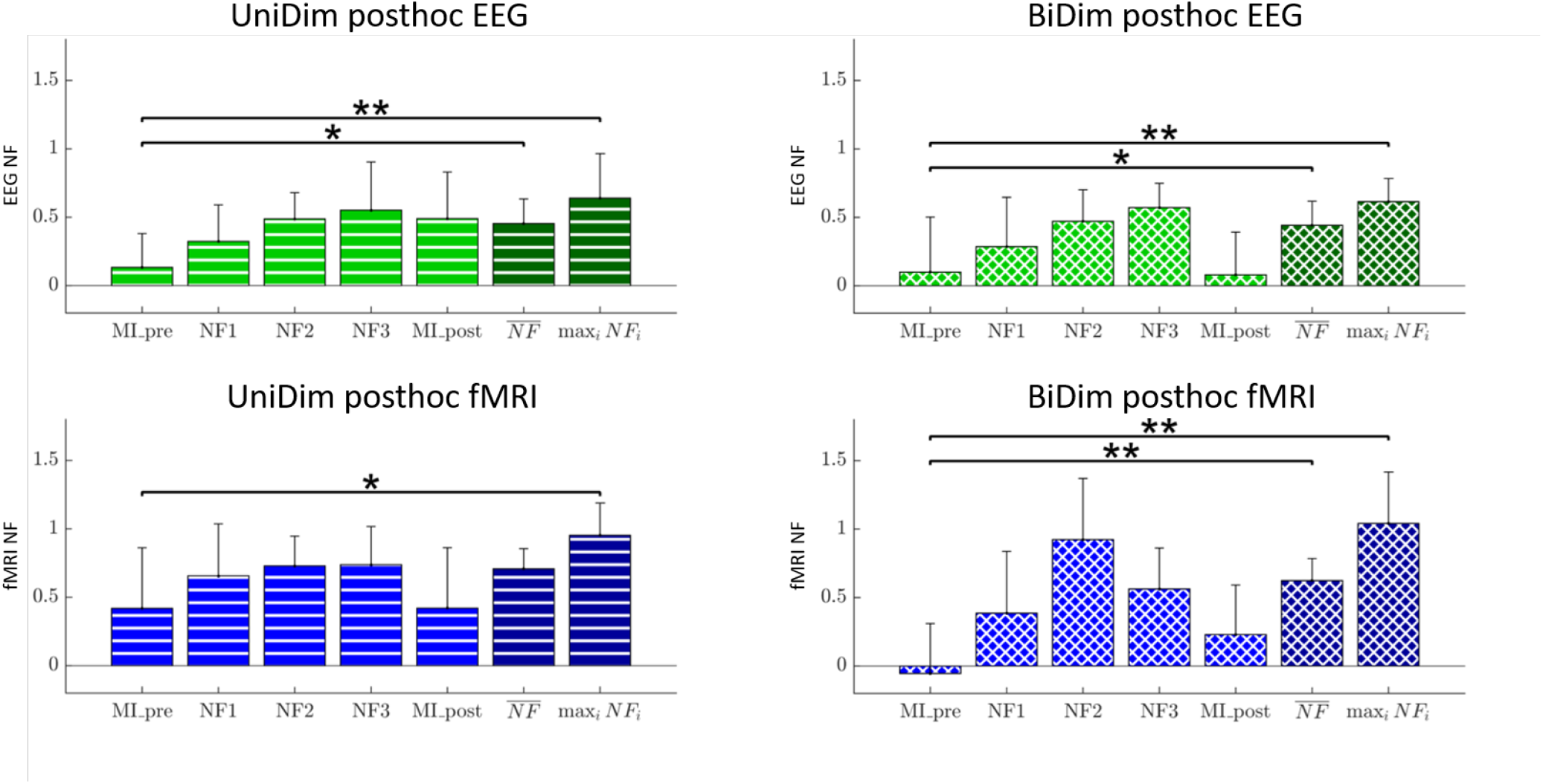
Group means (EEG/fMRI, posthoc, z-scored) on each run with standard deviation + significance of Wilcoxon tests.

Questionnaire results indicated that in the BiDim group 5 participants out of 10 found that the blocks were too short (against one who found them too long in the UniDim group), and 5 participants out of 10 found that the feedback was not a good indicator of their motor imagery (against 0 in the UniDim group).

## 4 DISCUSSION

In the present study we introduced and evaluated two integrated feedback strategies for EEG-fMRI-NF: a 2D metaphor in which EEG and fMRI are mapped onto each dimension, and a 1D gauge that integrates both information. In contrast to representing the EEG and fMRI features with two separate feedbacks, these integrated feedback strategies represent both information in a single feedback with a single NF target. The rationale behind the proposed integrated feedback was to reduce the cognitive load by training a single regulation task and at the same time feed back a richer NF information from both EEG and fMRI modalities.

### 4.1 Online and posthoc performance

Overall both strategies allowed participants to up-regulate MI-related EEG and fMRI patterns, as demonstrated by the higher posthoc EEG and fMRI activation levels during maxNF/NF compared to MI pre (see Fig 9). The improvement was even more significant on posthoc fMRI in the BiDim group.

Online fMRI activation level during NF were significantly higher in the UniDim group than in the BiDim group (Fig 8) which showed particularly high variability among participants and NF runs. Even if the UniDim worked better than the BiDim regarding the regulation of the initial (online) targets, their performance was moderate. Results suggests that the UniDim group had better online NF scores than the BiDim one. A significant improvement in online performances was only observed if the maximum NF scores were considered in the UniDim group. (see Fig 8). The loss of performance on the online fMRI activation level during NF with a bi-dimensional feedback was also observed in our previous study (Perronnet et al., 2017). Our new results thus highlight the fact that the bi-dimensional feedback is harder to control than the uni-dimensional feedback and that this affects online EEG and fMRI activation levels differently, at least on a single-session basis. We hypothesize that this could be due to the higher complexity of the bi-dimensional feedback. This complexity comes from the fact that it has two degrees of freedom with different update rates (4 Hz and 1 Hz), whose relationship is non-trivial, and one of which is delayed from the other. Subjects therefore need more time to familiarize with this more complex feedback. By allowing subjects to discriminate between the information coming from EEG and fMRI, they might be able to try different strategies and analyze how they affect both modalities. The separate regulation of EEG and fMRI modalities can be disturbing if they present inconsistencies. This could explain why half of the participants in the BiDim group reported that the feedback was not a good indicator of their motor imagery. The hypothesis that the bi-dimensional feedback is more complex and therefore requires more habituation time is supported by the fact that half of the participants in the BiDim group reported they found the training blocks too short (20 seconds) and by participants comments from the BiDim group: *”it is hard to know which mental process will favor EEG activity and which one will favor fMRI activity”, “the discrepancy between EEG and fMRI did not help to control the feedback given the small number of trials”, “task blocks could have been longer to allow to test different strategies and observe their effect”*. The fact that the loss of performance affected more fMRI than EEG could mean that they focused more on regulating the EEG because feedback from EEG is immediate while feedback from fMRI is delayed. Additionnally this could also be due to the fact that the feedback was moving 4 times faster in the EEG dimension.

Looking at the opposite trend between the online and posthoc activation levels of both groups (i.e. higher online fMRI activation levels for UniDim and higher posthoc fMRI activation elvels for BiDim) suggests that participants in the BiDim group could have moved further away from their initial MI pre calibration pattern than participants from the UniDim group. Even if the 2D feedback is more complex, it seems to encourage participants to explore mental strategies, interpret their effects on the two feedback dimensions in order to find a strategy that allows to control both dimensions equitably. Training block length might benefit from being adapted to the feedback strategy, with shorter block for the 1D feedback and longer block for the 2D feedback to allow for the exploration and interpretation of inner strategies. While the 2D strategy could prove valuable in the longer term to reach more specific self-regulation, the 1D metaphor could be well suited during earlier phases of a NF program or for clinical application as it is easier to control. This is also the reason why we opted for the 1D metaphor in a pilot study where we tested the feasibility of EEG-fMRI NF for stroke rehabilitation (Lioi et al., 2020).

### 4.2 Group distribution across the 3 NF runs

Looking at the distribution of online mean activation levels (Fig 10) over the three NF runs shows how the two group populations evolved over the course of the training. In the first run, both populations were rather widespread and distributed along the EEG axis which suggests that participants started by exploring EEG. Participants from the BiDim group were also slightly distributed along the fMRI axis in the first NF run. In the second NF run, both populations were spread along the fMRI axis, which suggests that participants explored fMRI while keeping EEG at a mean level. In the third run, both populations are spread along the central (0.5) isoline, which suggests that participants adopted a strategy that minimized the errors in both dimensions. Overall the progression look similar in both group but the BiDim population is more widespread than UniDim in NF1 and NF2. This higher variability might once again be due to the higher complexity of the feedback to which participants need to get used to.

**Figure 10.**
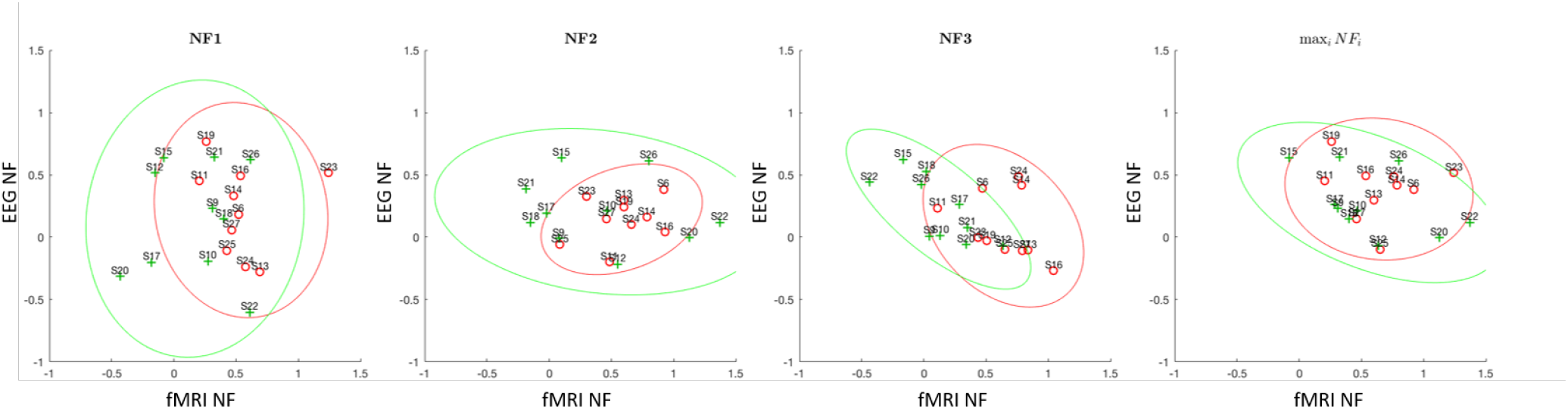
Individual means (online EEG NF and fMRI NF, z-scored) of all participants during NF runs. Individuals from the UniDim group are shown in red. Individuals from the BiDim group are shown in green.

**Figure 11.**
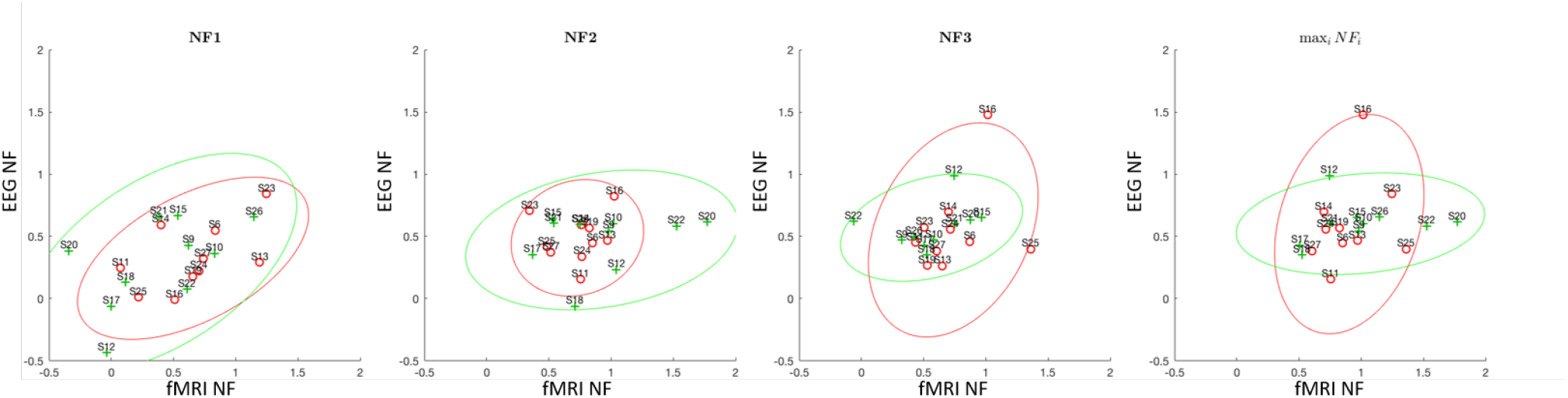
Individual means (posthoc EEG NF and fMRI NF, z-scored) of all participants during NF runs. Individuals from the UniDim group are shown in red. Individuals from the BiDim group are shown in green.

Accurate assessments of strategies, motivation, cognitive state but also of personality traits (Jeunet et al., 2015a, 2018) are key to interpret the learning curve and the effects of NF training. Our questionnaire inquiring about strategies, fatigue, motivation, perceived performance, and confidence in the feedback gave us valuable qualitative knowledge about how subjects in the BiDim group reacted to the complexity of the feedback. In future experiments, especially some involving multiple NF sessions, we plan to use a more thorough questionnaire incorporating behavioural and personality aspects.

### 4.3 Activation maps

BOLD activation maps show that during NF both groups significantly activated regions from the motor imagery network including premotor areas and posterior parietal areas (figures 5 and 4), as well as regions that have been shown to be consistently active during NF (Emmert et al., 2016) (mid-cingulate, supra-marginal cortex (temporo-parietal), dlPFC, premotor cortex). Subcortical and cerebellar regions activations could not be identified as they were out of the field of view. The BiDim group showed more activations (p<0.001, uncorrected) than the UniDim group in the right superior parietal lobule (BA 7). The SPL plays an essential role in many cognitive, perceptive, and motor-related processes (Wang et al., 2015; Culham and Kanwisher, 2001). In particular it has been reported to be activated both during motor execution and MI (Solodkin et al., 2004; Raffin et al., 2012; Lotze and Halsband, 2006; Hétu et al., 2013; Confalonieri et al., 2012; Sharma and Baron, 2013; Fleming et al., 2010) with greater activation has been observed during imagined rather than executed movement (Gerardin et al., 2000; Hanakawa et al., 2002). More specifically, the SPL is known to play a role in guiding motor activity in relation to spatial information (Buneo and Andersen, 2006; Wang et al., 2015; Culham and Kanwisher, 2001) and to be crucial in the generation of mental motor representations (Sirigu et al., 1996). Several studies have demonstrated that impairments to the parietal cortex reduced MI ability (Sirigu et al., 1996; Danckert et al., 2002; McInnes et al., 2016). A recent meta-analysis showed that patients with parietal lobe damage exhibited most impaired MI ability (McInnes et al., 2016). In MI, the SPL is thought to play a role in facilitating the planning and coordination of imagined movements and/or in indirectly inhibiting M1 through its connection with the SMA (McInnes et al., 2016; Kasess et al., 2008; Solodkin et al., 2004). Activations in the SPL have been shown to be more active during visual imagery than during kinaesthetic imagery (Guillot et al., 2009). However we found no significant higher activation in the occipital regions in BiDim as would be expected during visual imagery. Therefore it is unlikely that the SPL activation would indicate that participants in the BiDim performed a motor imagery that would have been more visual than kinesthetic. The superior parietal cortex has also been demonstrated to be active when feedback is presented visually (Sitaram et al., 2016; Emmert et al., 2016; Ninaus et al., 2013). However the fact that the SPL was more significantly active in the BiDim group than in the UniDim group suggest that it is more than a generalized NF effect. This activation could result from both the overlap of the motor imagery task and the self-regulation process (Sitaram et al., 2016), both of which could be more intense under the bi-dimensional condition.

The overlay of UniDim and BiDim fMRI activations (see Fig 5) shows that activations in the premotor areas were more widespread and bilateral in the UniDim group while they were more localized and lateralized to the left hemisphere in the BiDim group. Also, the BiDim group showed significant deactivations in the right primary motor cortex while the UniDim group did not. Overall, our results suggest that the bi-dimensional feedback triggered more specific activations than the uni-dimensional feedback.

### 4.4 Defining bimodal NF targets

An integrated feedback allows to reward specific EEG/fMRI features and gives flexibility on the definition of the bimodal NF target, depending on the assumed spatio-temporal complementarity of the EEG and fMRI features. In this study, we designed integrated feedback so that subjects would have to regulate both EEG and fMRI at the same time in order to reach the NF target. This assumes that such a state is possible. Indeed, neuro-vascular studies show that the electrophysiological and hemodynamic activity are correlated (Formaggio et al., 2010; Gonçalves et al., 2006; Ritter et al., 2009; Zaidi et al., 2015; Murta et al., 2015). For example, a study by Zaidi et al. (Zaidi et al., 2015) found significant correlations between hemodynamic peak-times of oxygenated (HbO2) and deoxygenated (HbR) hemoglobin signals with the underlying neural activity measured with intra-cortical electrophysiology in primates. However depending on the type of tasks, the features, and the subjects, this might not necessarily be the case as illustrated in the study by De Vos et al. (De Vos et al., 2013) who reported no correlation between EEG and fMRI of a face processing task. Even if it is challenging to predict the degree of complementarity and redundancy of the EEG and fMRI features, it might be beneficial to take into consideration the degree of correlation of both features during the calibration phase. However a preliminary study (Cury et al., 2020) proposed a model able to learn, despite noise of simultaneous acquisition, from one modality (fMRI) to predict neurofeedback scores on the other (EEG), showing a certain degree of coherence between the two. The study used the data presented in this paper.

Instead of defining the target on the “intersection” of the EEG and fMRI features, one could think of using an easier target defined by their “union” (the target would be reached when the EEG target or the fMRI target is reached). Such a target would be easier to reach, therefore potentially less specific, but it might be advantageous in order to limit the user frustration when used at the beginning of a protocol for example. Also the “union” strategy could be used in case the EEG and fMRI features would be hardly redundant. This could happen if the mental process being regulated was more complex and involved for example a cognitive regulation and an emotional regulation aspect each of which would be associated to one of the feature. Moreover in the “union” strategy, one could imagine displaying a secondary reward when the pair of EEG and fMRI features would reach the intersection without “penalizing” the subject when he/she does not control for both.

### 4.5 Limitations

The strong artifacts affecting the EEG in the MR environment and the need to correct them in real-time constitute the main current limitation of EEG-fMRI-NF. Artifact removal for simultaneous EEG/fMRI traditionnally consists of correcting the gradient and the BCG artifact with the average artifact subtraction technique. However, this technique is likely to result in residual artifacts. In their pionneering work, Zotev et al. reported that residual MR and CB artifacts contributed up to 50% to the EEG feature after basic real-time signal processing (Zotev et al., 2014). Other sources of artifacts such as the helium pump (Nierhaus et al., 2013), the MR ventilation and motion (Jansen et al., 2012; Fellner et al., 2016) can also seriously affect the EEG data. As the helium pump artifact lies in the gamma range, in our particular case it should not affect our features of interest (alpha/beta). But altogether, these other sources of artifacts can limit the quality of the online EEG-NF signal and therefore also the potential of the EEG-fMRI-NF approach. However, the posthoc CSP patterns of the majority of the participants corresponded to motor imagery patterns which confirms that participants executed the task well and that the data was well pre-processed.

Alternative to the average artifact subtraction procedure such as optimal basis sets (Niazy et al., 2005), independent component analysis (Mantini et al., 2007), reference-layer adaptive subtraction (Chowdhury et al., 2014) and carbon-wire loop (van der Meer et al., 2016; Abbott et al., 2014) have been proposed in order to better remove gradient and BCG artifacts or to remove the other types of artefacts. However only few methods are currently available for online use (Krishnaswamy et al., 2016; Mayeli et al., 2016; Wu et al., 2016; van der Meer et al., 2016; Steyrl et al., 2017). In future experiments, using such methods could allow to improve the EEG-NF signal quality. Interestingly, a recent approach called automated EEG-assisted retrospective motion correction (aE-REMCOR) (Wong et al., 2016)) uses the EEG data in order to estimate head motion and improve fMRI motion correction. Another limitation of this study is that we did not monitor movements of the hand during the motor imagery task by measuring the electromyographic signal. This measure requires the installation of an additional amplifier and needs custom cable lengths for each individual therefore increasing the burden and complexity of the simultaneous EEG-fMRI setup. Upper limb movements were therefore monitored by means of a camera inside the MR bore and subjects were repeatedly instructed to avoid movements.

## 5 DATA AVAILABILITY

We have uploaded the anonymized EEG and fMRI datasets this study in BIDs format in a open-access repository https://openneuro.org/datasets/ds002338 to promote reuse. We have complemented the raw data with a series of derivatives: NF EEG and fMRI scores and head motion time-series for each subject and training run. We also calculated the framewise displacement as in (Power et al., 2014) and performed a correlation analysis with EEG and fMRI NF scores to assess the impact of movements artifacts. A detailed description of the dataset architecture and metadata can be found in (Lioi et al., 2019).

## 6 CONCLUSION

Our study introduces new integrated feedback strategies for EEG-fMRI-NF and demonstrates that during a motor imagery task they enable to regulate EEG and fMRI simultaneously, even when EEG and fMRI are integrated in a 1D feedback. Our results also suggest that the 1D feedback is easier to control on a single session while the 2D feedback encourages subjects to explore their strategies to find one that allows to control EEG and fMRI by recruiting more specific brain patterns. Altogether, our study paves the way to novel integrated EEG-fMRI-NF strategies for the development of flexible and effective NF paradigms that could prove useful for clinical applications.

## 7 ACKNOWLEDGEMENTS

This work has received a French government support granted to the CominLabs excellence laboratory and managed by the National Research Agency in the “Investing for the Future” program under reference ANR-10-LABX-07-01. It was also financed by Brittany region under HEMISFER project, and the National Research Agency with the REBEL project and grant ANR-15-CE23-0013-01. MRI data acquisition was supported by the Neurinfo MRI research facility from the University of Rennes I. Neurinfo is granted by the European Union (FEDER), the French State, the Brittany Council, Rennes. F.L. was supported by the European Research Council with project BrainConquest (grant ERC-2016-STG-714567). We thank Jean Alla for proofreading the paper.

## Notes

#### Summary of Updates

correction of phrasing and typos to improve clarity

